# Ocellar spatial vision in *Myrmecia* ants

**DOI:** 10.1101/2021.05.28.446117

**Authors:** Bhavana Penmetcha, Yuri Ogawa, Laura A Ryan, Nathan S Hart, Ajay Narendra

## Abstract

In addition to the compound eyes insects possess simple eyes known as ocelli. Input from the ocelli modulates optomotor responses, flight-time initiation and phototactic responses, behaviours that are predominantly mediated by the compound eyes. In this study, using pattern electroretinography (pERG), we investigated the contribution of the compound eyes to ocellar spatial vision in the diurnal Australian bull ant, *Myrmecia tarsata* by measuring the contrast sensitivity and spatial resolving power of the ocellar second-order neurons under various occlusion conditions. Furthermore, in four species of *Myrmecia* ants active at different times of the day and in European honeybee, *Apis mellifera*, we characterized the ocellar visual properties when both visual systems were available. Among the ants, we found that the time of activity had no significant effect on ocellar spatial vision. Comparing day-active ants and the honeybee we did not find any significant effect of locomotion on ocellar spatial vision. In *M. tarsata*, when the compound eyes were occluded, the amplitude of the pERG signal from the ocelli reduced by three times compared to conditions when the compound eyes were available. The signals from the compound eyes maintained the maximum contrast sensitivity of the ocelli as 13 (7.7%), and the spatial resolving power as 0.29 cpd. We conclude that ocellar spatial vison improves significantly with input from the compound eyes, with a noticeably larger improvement in contrast sensitivity than in spatial resolving power.

## Introduction

Insects use their visual systems to detect relevant information from their environment to orient themselves, find conspecifics, forage, navigate, hunt and mate (Cronin et al., 2014). While the compound eyes have been extensively studied (eg., Greiner, 2006; Land, 1989; Narendra et al., 2017; Warrant, 2008), many insects also possess a single-lens eye known as an ocellus which has been relatively understudied (eg., Mizunami, 1995; Ribi and Zeil, 2018; Warrant et al., 2006). Typically, three simple eyes are placed in a triangular formation on the dorsal surface of the head. Each ocellus consists of a lens, an iris, a corneageous cell layer and a retina differentiated into dorsal and ventral retinae (Narendra and Ribi, 2017; Ribi and Zeil, 2018; Ribi et al., 2011). Almost all flying insects possess ocelli. Their functions have been best studied in dragonflies and locusts where the ocelli play a crucial role in horizon detection (Berry et al., 2007a; Stange et al., 2002) and attitude control during flight (Mizunami, 1995; Stange, 1981; Taylor, 1981; Wilson, 1978). In addition, input from the ocelli aids visually guided behaviours such as flight time initiation (Eaton et al., 1983; Lindauer and Schricker, 1963; Schricker, 1965; Wellington, 1974), optomotor responses (Honkanen et al., 2018) and phototactic responses (Barry and Jander, 1968; Cornwell, 1955). Ocelli are typically absent in walking insects. Exceptions to this include workers of the desert ants of the genus *Cataglyphis* and *Melophorus* (Penmetcha et al., 2019; Schwarz et al., 2011a) and bull ants or jack jumpers of the genus *Myrmecia* (Narendra and Ribi, 2017; Narendra et al., 2011). Behavioural evidence shows that the ocelli of the desert ants derive compass information from celestial cues (Schwarz et al., 2011b; Schwarz et al., 2011c), especially the pattern of polarized skylight (Fent and Wehner, 1985; Mote and Wehner, 1980).

Neuroanatomical studies have confirmed interactions between the ocelli and the compound eye specifically between the large second-order neurons, L-neurons, that receive input from a large number of ocellar photoreceptors, and the optic lobe where signals from the compound eyes are processed. For example, in honeybees and flies, efferent fibres run from the lobula plate into the ocellar tract (Strausfeld 1976, see also Hagberg and Nässel 1986). In the European field cricket *Gryllus campestris*, fibres run from the medulla to the ocellar tract forming knob-like blebs (Honegger and Schürmann, 1975). In the Australian field cricket, *Teleogryllus commodus*, and house cricket, *Acheta domesticus*, neurons extend from the ocellar photoreceptors to the lobular layers of the compound eyes and additionally in the laminar layers of *T. commodus* (Rence et al., 1988). Physiological studies in *T. commodus* showed that the amplitude of the electroretinograms (ERGs) measured from the compound eyes was reduced by 20% following ocellar occlusion (Rence et al., 1988). However, it is unknown whether the visual information received by the compound eyes has an effect on the visual capabilities of the ocelli.

The capabilities of a visual system is determined by the extent to which it can discriminate between fine details of objects in a scene (spatial resolving power) and adjacent stimuli based on differences in relative luminosity (contrast sensitivity) (Land, 1997). The image quality of ocellar lenses have been estimated histologically, using the hanging drop technique (originally described by Homann, 1924) and modifications of this technique. Histological evidence suggests that in some insects the ocellar lenses likely produce under-focused images because the focal plane is located behind the retina in migratory locusts, *Locusta migratoria* (Berry et al., 2007b; Wilson, 1978), sweat bees, *Megalopta genalis* (Warrant et al., 2006), blowflies, *Calliphora erythrocephala* (Cornwell, 1955; Schuppe and Hengstenberg, 1993), orchid bees, *Euglossa imperialis* (Taylor et al., 2016) and Indian carpenter bees, *Xylocopa leucothorax, X. tenuiscapa, X. tranquebarica* (Somanathan et al., 2009). In some insects the ocelli appear to be capable of resolving images with the plane of best focus located close to the retina as seen in European honeybees, *Apis mellifera* (Ribi et al. 2011, but see Hung and Ibbotson 2014), paper wasps, *Apoica pallens, Polistes occidentalis* (Warrant et al., 2006) and dragonflies, *Hemicordulia tau, Aeshna mixta* (Berry et al., 2007c; Berry et al., 2007a; Stange et al., 2002). Additionally, the spatial resolution of the honeybee ocelli was estimated by quantifying the contrast in the image produced by the lens (Hung and Ibbotson, 2014) and that of the dragonfly ocelli by measuring the acceptance angles of the ocellar photoreceptors (Berry et al., 2007c). In honeybees, when provided with vertical and horizontal gratings, it was found that contrast information for high spatial frequency was transferred through the ocellar lenses better than low spatial frequency (Hung and Ibbotson, 2014). In dragonflies, the acceptance angles were obtained by ray tracing through anatomical models of the median ocellar lens and retina. The acceptance angles were two times lower in elevation (5.2°) than in azimuth (10.3°) suggesting higher resolution in the vertical plane (Berry et al., 2007c).

The hanging drop technique and other histological methods do not consider the physiological properties of the photoreceptors or the ocellar second-order neurons which is essential to determine the visual capabilities of the eye. Hence, intracellular electrophysiology has been used to infer the spatial resolution of the ocelli. In dragonflies, the spatial resolution of the ocelli was extrapolated by measuring the angular sensitivities of the median ocelli photoreceptors (van Kleef et al., 2005) and ocellar second-order neurons (Berry et al., 2006; Berry et al., 2007a) in response to green and ultra-violet (UV) LED arrays. Similar experiments were done on the second-order neurons in the lateral ocelli of locusts (Wilson, 1978) in response to a Xenon arc lamp. In dragonfly median ocelli, the acceptance angles of the photoreceptors were 15° in elevation and 28° in the azimuth, in the vertical and horizontal plane respectively indicating relatively enhanced spatial resolution in the vertical plane (van Kleef et al., 2005). These values were a factor of 2 or more larger than when obtained from the ray tracing method mentioned previously (Berry et al., 2007c). Additionally, although the acceptance angles were slightly higher (elevation = 20°, azimuth = 40°) when measured from the ocellar second-order neurons, the hypothesis of enhanced spatial resolution in the vertical plane remains unchanged; providing evidence that spatial resolution is conserved after convergence of photoreceptors onto second-order neurons (Berry et al., 2006). In locusts, angular sensitivities were measured only in the horizontal plane and it was found that the field widths measured at 50% maximum sensitivity showed considerable variation. However, the total extent of the field for all the cells showed less variation and was at least 130° indicating that the spatial resolution was low in these neurons (Wilson, 1978). Therefore, while the spatial resolution of the ocelli has been estimated using various techniques, the contrast sensitivity of the ocelli has neither been estimated nor measured physiologically (but see Simmons, 1993 for intracellular responses of L-neurons to sinusoidally modulated light of varying contrasts in locusts).

Pattern electroretinography (pERG) has been established to simultaneously measure spatial resolving power and contrast sensitivity from the compound eyes in ants (Ogawa et al., 2019; Palavalli-Nettimi et al., 2019) and honeybees (Ryan et al., 2020). The pERG consists of non-linear ERG components that are generated by second-order visual neurons at least in vertebrates (Porciatti, 2007) in response to contrast-reversing sinusoidal gratings that are contrast-modulated patterned visual stimuli at constant mean luminance. In insects, when measured in the presence of ON and OFF light stimuli, ERGs from the compound eyes show the presence of ON transients (Coombe, 1986; Ryan et al., 2020) and OFF transients (Coombe, 1986; Palavalli-Nettimi et al., 2019; Ryan et al., 2020) arising from the second-order neurons in the lamina. ERGs from the ocelli of some insects show the presence of Component 3, a hyperpolarizing post-synaptic potential and a sustained after-potential, arising from the ocellar second-order neurons (Ruck, 1961a; Ruck, 1961b). Hence, the pERG technique is ideal as it allows us to simultaneously determine the spatial resolving power and contrast sensitivity.

Ants of the genus *Myrmecia* are unusual, as closely related congeneric species are active at different times of the day (Greiner et al., 2007; Narendra et al., 2011). Each species has evolved distinct adaptations in their compound eyes (Greiner et al., 2007; Narendra et al., 2011; Narendra et al., 2017; Ogawa et al., 2019) and ocelli (Narendra and Ribi, 2017; Narendra et al., 2011) to suit the specific temporal niches they occupy. The ocelli of night-active *Myrmecia* ants tend to have larger lenses and wider rhabdoms to improve optical sensitivity (Narendra and Ribi, 2017). Among worker ants, *Myrmecia* have relatively large ocelli which makes this group ideal to investigate ocellar physiology in day- and night-active ants.

In this study we determined the contribution of the compound eyes to ocellar spatial vision using the pERG technique. We measured contrast sensitivity and spatial resolving power of the ocellar second-order neurons in the diurnal-crepuscular *Myrmecia tarsata* under different visual system occlusion conditions. Next, we measured the ocellar visual properties in four *Myrmecia* species to explore the effect of activity time on ocellar spatial vision. Lastly, to identify the effect of locomotory style on spatial vision we compared the ocellar visual properties of diurnal walking *Myrmecia* ants to the diurnal flying honeybee, *Apis mellifera*.

## Methods

### Study Species

We studied the physiology of the median ocelli in workers of four *Myrmecia* ant species: diurnal-crepuscular *Myrmecia gulosa* (Sheehan et al., 2019) and *Myrmecia tarsata* (Greiner et al., 2007); and the strictly nocturnal *Myrmecia midas* (Freas et al., 2017) and *Myrmecia pyriformis* (Greiner et al., 2007; Narendra et al., 2010). To identify whether ocellar spatial vision is influenced by locomotory differences, we compared the spatial properties of the day-active worker ants that have a pedestrian lifestyle with that of the workers of the day-active European honeybee *Apis mellifera* that predominantly fly. The animals were captured from multiple nests at the following locations: *M. midas, M. tarsata* and *A. mellifera* from Macquarie University campus, Sydney, NSW (33°46’10.24”S, 151°06’39.55”E); *M. gulosa* from Western Sydney University campus, Hawkesbury, Sydney, NSW (33°37’46.35”S, 150°46’04.47”E); *M. pyriformis* from The Australian National University campus, Canberra, ACT (35°16’50”S, 149°06’43”E).

### Animal preparation

To perform electrophysiological recordings, the ants were anesthetized by cooling them in an icebox for 5-10 mins before removing their antennae, legs and gaster. Each animal was then fixed horizontally onto a plastic stage with its dorsal side facing upwards using beeswax. The orientation of the median ocelli varies slightly between species (Narendra and Ribi, 2017). For instance, in *M. midas* and *M. pyriformis* the median ocelli are upward-facing, whereas in *M. tarsata* and *M. gulosa* the median ocellus are forward-facing. Hence, in our preparations we oriented the head to ensure that the median ocellus faced the stimulus. To place an active electrode on the median ocellar retina, a small incision was made using a sharp blade and the cuticle was thinned immediately posterior to the median ocellar lens. Vaseline (Unilever, USA) was placed on the thinned cuticle to prevent dehydration and a layer of conductive gel (Livingstone International Pty Ltd., New South Wales, Australia) was added. The honeybees were prepared similar to the ants after anesthetizing, following which their wings and legs were removed. The hair around the median ocellus was removed using sharp forceps for easier access to the retina and placement of electrode. In the honeybees, incisions were not required as their cuticle was thinner than the ants.

The animals were then mounted within a Faraday cage wherein electrophysiological recordings were carried out on the median ocellar retinae. An active electrode of platinum wire of 0.25mm diameter with a sharp tip was placed at the point where the cuticle was thinned, posterior to the median ocellar lens in the ants. In the honeybees the active electrode was placed on the cuticle, posterior to the median ocellar lens to access the retina underneath. The active electrode was immersed in the conductive gel in both cases. A silver/silver-chloride wire of 0.1mm diameter was inserted into the mesosoma of the ants and the thorax of the honeybees which served as an indifferent electrode.

To reduce any effects of the circadian rhythms on eye physiology, the experiments were conducted at the activity time of each species, i.e., from 1-6 hours post-sunset for nocturnal species and 2-8 hours post-sunrise for the diurnal species.

### Electroretinogram

We measured the ON-OFF responses using electroretinography from the median ocelli in *M. tarsata* (n=5) to confirm the presence of extracellular potentials produced by the ocellar second-order neurons in electroretinograms (ERGs).

A white LED light source (5mm in diameter with irradiance of 5.81 × 10^−5^ W cm^-2^, C503C-WAS-CBADA151, Cree Inc., Durham, NC, USA) was used as a stimulus and placed 15 cm from the ocelli. The ants were dark adapted for 5 mins prior to stimulation. Ten consecutive trials were carried out, each trial lasting for 10 seconds with the ON-OFF LED light stimulus presented after an initial delay of 0.5 seconds and for a total ON duration of 5 seconds using a custom-written script in MATLAB (R2015b, Mathworks, Natick, MA, US). The signals were filtered between 0.1 Hz – 100 Hz and amplified at x100 gain (AC) using a differential amplifier (DAM50, World Precision Instruments Inc., FL, USA).

Certain ocellar second-order neurons (L-neurons) are known to form synapses with the descending interneurons which receive input from the compound eyes ((Strausfeld, 1976; *Apis mellifera* (Guy et al., 1979); *Schistocera gregaria, Schistocera americana* (Rowell and Pearson, 1983); *Calliphora erythrocephala* (Strausfeld and Bassemir, 1985)). Hence, we measured the ERGs from the median ocelli retina of *M. tarsata* for individuals with both visual systems-intact (E^+^O^+^, n=5), and again with their compound eyes-occluded (E^-^O^+^, n=5). These two experiments were conducted on the same individual consecutively in random order. Occlusion was done using black nail enamel (B Beauty, NSW, Australia) and could be completely peeled off without damaging the eye.

### Pattern electroretinogram (pERG)

We used pattern electroretinography to measure the spatial resolving power and the contrast sensitivity of the ocelli of *Myrmecia* ants and *Apis mellifera* (n=4 for *M. gulosa*, n=5 for all other species).

The pERG visual stimuli were projected by a digital light processing projector (W1210ST, BenQ corporation, Taipei, Taiwan) onto a white screen (W51 x H81cm) at a distance of 30 cm from the animal. The stimuli were vertical contrast-reversal sinusoidal gratings of different spatial frequencies (cpd: cycles per degree) and Michelson’s contrasts. They were generated using Psychtoolbox 3 (Pelli, 1997) and MATLAB (R2015b, Mathworks, Natick, MA, US) and controlled via a custom Visual Basic Software written in Visual Studio (2013, Microsoft Corporation, Redmond, WA, US). The mean irradiance of the grating stimulus was 1.75 × 10^−4^ W/cm^2^ which was measured using a calibrated radiometer (ILT1700, International Light Technologies, Peabody, MA, US). A temporal frequency of 4 Hz was used for all the stimuli.

Prior to the first recording the animal was adapted to a uniform grey stimulus with the same mean irradiance as the grating stimuli for 20 minutes. To measure the contrast sensitivity function of the ocelli, nine spatial frequencies (0.05, 0.1, 0.15, 0.2, 0.26, 0.31, 0.36, 0.41, 0.52 cpd) and five contrasts (95%, 50%, 25%, 12.5%, 6%) for each spatial frequency were presented. The spatial frequencies were first presented in decreasing order of frequencies (0.52, 0.36, 0.26, 0.15, 0.05 cpd), skipping one frequency in between. In order to evaluate any degradation in response over time the interleaved frequencies were then presented in ascending order (0.1, 0.2, 0.31, 0.41 cpd). At each spatial frequency, all five contrasts were tested in decreasing order. Each combination of the stimuli was recorded for fifteen repeats, five seconds each and averaged in the time domain. The averaged responses were then analysed using a Fast Fourier Transform (FFT) in the frequency domain. As a control, the non-visual electrical signal (background noise) was recorded at two spatial frequencies (0.05 and 0.1 cpd) at 95% contrast with a black board to shield the animal from the visual stimuli before and after running the experimental series. The maximum signal out of four control runs was used as the noise threshold.

### Contribution of the two visual systems to the pERG response amplitude in M. tarsata

To identify the origin of the neural responses in pERG, we compared the pERG response amplitudes in *M. tarsata* measured at each of the nine spatial frequencies (at 95% contrast) under different conditions: (a) both visual systems-intact (E^+^O^+^) (n=5), (b) ocelli-occluded (E^+^O^-^) (n=3), (c) compound eyes-occluded (E^-^O^+^) (n=4) and (d) visual systems-intact with incisions in the lamina (E^+^O^+^L^-^) (n=3). In condition d, we made a rectangular window on the dorso-frontal region of both compound eyes using a sharp blade. We then accessed the lamina and made incisions on the basement membrane on both sides of the retinae using a sharp blade. The windows were subsequently covered with vaseline to prevent dehydration and the recordings were carried out. Occlusions were done as mentioned above.

### Data analysis

#### Spatial resolving power and contrast threshold

As the spatial frequency of the visual stimuli increases the amplitude of the pERG response at the second harmonic (8Hz) decreases. The point at which the pERG response amplitude dropped below the noise threshold was defined as the spatial resolving power at the highest contrast. At each spatial frequency of the visual stimuli, the amplitude of the pERG response decreased with decreasing contrast. The point at which the amplitude dropped below the noise threshold was taken as the contrast threshold. The contrast threshold at each spatial frequency of gratings was used to calculate the contrast sensitivity at that particular spatial frequency using the formula: 1/contrast threshold. (See Ogawa et al., 2019 for more details)

### Statistical Analysis

We used a linear-mixed effects model in RStudio (R Core Team, 2018) to test whether the spatial frequency of the stimulus and the treatment conditions (E^+^O^+^, E^+^O^-^, E^-^O^+^, E^+^O^+^L^-^) had an effect on the pERG response amplitudes in *M. tarsata*. Spatial frequency of the stimulus and the treatment conditions were the fixed effects and animal identity was a random effect. The significances of the fixed effect terms were examined using Type III ANOVA. Both pERG response amplitudes and the spatial frequency data were log-transformed before the analysis and the final residuals were plotted and met the model assumptions.

To assess the effect of treatment condition (E^+^O^+^, E^+^O^-^) and spatial frequency of stimulus on the contrast sensitivity function in *M. tarsata* we used a linear-mixed effects model in R. The condition and the spatial frequency were the fixed effects and animal identity was a random effect. Using a linear model, we tested whether the spatial resolving power and the maximum contrast sensitivity differed between the two conditions. The contrast sensitivity and spatial frequency data were log-transformed before the data analysis. Final residuals of the data were plotted and met the model assumptions.

Lastly, we used a linear model to test whether the maximum contrast sensitivities differed between the four *Myrmecia* species, and the spatial resolving powers differed among the four *Myrmecia* species and *Apis mellifera*. A linear-mixed effects model was used to assess the effect of species, their time of activity and spatial frequency of stimulus on the contrast sensitivity function of the four *Myrmecia* species using a maximum likelihood (ML) estimation method. The same was done, among diurnal *Myrmecia* ants and *Apis mellifera*, to assess the effect of the species and their mode of locomotion on the contrast sensitivity functions. Time of activity in ants, spatial frequency and locomotion were used as fixed effects and animal identity nested within species was used as a random effect. The significances of the fixed effect terms were examined using the t-test with Satterthwaite approximation for degree of freedom (ImerTest package). The model also reflected the variability in the dependent variable (contrast sensitivity function) due to the random effects. The contrast sensitivity and spatial frequency data were log-transformed before the data analysis and the final residuals of the data were plotted and met the model assumptions. All linear-mixed effect models were carried out in the *lme4* package of R (https://cran.r-project.org/web/packages/lme4/index.html).

## Results

### Electroretinogram

We first recorded ERGs to ON-OFF LED stimuli from the median ocelli of *M. tarsata* under two treatment conditions: E^+^O^+^ (pink line in Fig. 1) and E^-^O^+^ (black line in Fig. 1) to confirm the presence of the extracellular potentials originating from the ocellar second-order neurons. In E^+^O^+^ condition (pink line in Fig. 1B, C) we identified at the stimulus onset, Component 3, a hyperpolarizing post-synaptic potential (arrow in Fig. 1B) and the sustained after-potential of Component 3 was also identified at the stimulus offset (arrow in Fig. 1C). However, in E^-^ O^+^ treatment condition (black line in Fig. 1B, C) Component 3 was identified at the stimulus offset (arrow in Fig. 1C) and was less pronounced at the stimulus onset (dashed arrow Fig.1B).

**Figure 1.**
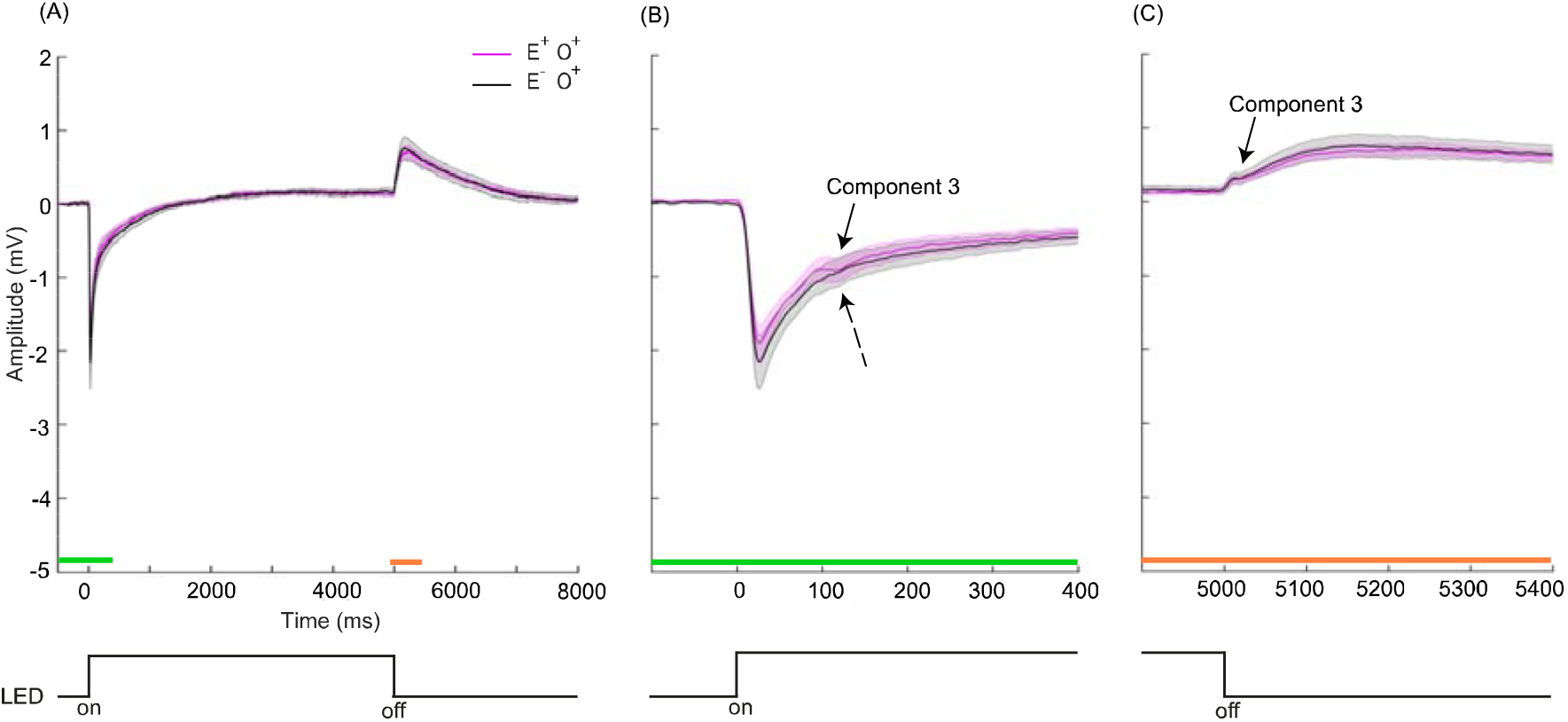
Electroretinograms from median ocelli of *Myrmecia tarsata* for visual systems-intact (E^+^O^+^) and compound eyes-occluded (E^-^O^+^) individuals. Mean ERGs from ocelli in response to ON-OFF LED light stimulus for visual systems-intact (E^+^O^+^, n=5) individuals are shown in pink and for that of compound eyes-occluded (E^-^O^+^, n=5) individuals are shown in black. The shaded regions show the standard error for the respective conditions. Solid arrows indicate presence of Component 3. Dashed arrow shows that Component 3 is less pronounced in E^-^O^+^ treated individuals. Stimulus onset and offset represented at the bottom. **(A)** ERGs for E^+^O^+^ and E^-^O^+^ treated individuals. Green bar indicates time scale magnified and presented in **(B)** and orange bar indicates time scale magnified and presented in **(C)**.

### Contribution of the two visual systems to the pERG response amplitude in M. tarsata

We measured the pERG response amplitudes from the ocellar second-order neurons in *M. tarsata* for each spatial frequency at 95% contrast. To confirm whether the ocellar second-order neurons receive inputs from the compound eyes in addition to the ocellar photoreceptors, we compared the response amplitudes in E^+^O^+^, E^-^O^+^, E^+^O^-^ and E^+^O^+^L^-^ individuals (Fig. 2A, B, C, D). The spatial frequency of the visual stimuli and the treatment conditions had a significant effect on the response amplitudes (Linear-mixed effect model, parameter=treatment condition, df=3, F=3.31, *P*=0.03; parameter=spatial frequencies, df=1, F=141.23, *P*<2.2e-16; parameter=treatment condition:spatial frequency, df=3, F=19.51, *P*=4.06e-10). For E^+^O^+^, E^-^O^+^ and E^+^O^-^ treated individuals, the response amplitudes decreased with increasing spatial frequencies (slope for E^+^O^+^= −1.06, E^-^O^+^= −0.62, E^+^O^-^= −1.32, E^+^O^+^L^-^ = 0.02) (Fig. 2A, B, C). The response amplitudes for E^+^O^+^ individuals were significantly different from the E^+^O^+^L^-^ (Linear-mixed effect model, *t*=6.24, *P*=9.5e-09) and E^-^O^+^ (*t*=2.7, *P*=0.008) treated individuals. Similarly, response amplitudes for E^+^O^-^ treated individuals were significantly different from E^+^O^+^L^-^ (*t*=7.1, *P*=2.34e-10) and E^-^O^+^ (*t*=3.85, *P*=0.0002) treated individuals. Additionally, the response amplitudes were significantly different between E^+^O^+^L^-^ and E^-^O^+^ (*t*=-3.37, *P*=0.001) treated individuals but not significantly different between E^+^O^+^ and E^+^O^-^ (*t*=-1.59, *P*=0.12) treated individuals. At the lowest spatial frequency (0.05cpd) the mean pERG response amplitude for E^+^O^+^ was 0.009 ± 0.001 (mean ± S.E.) mV (Fig. 2A), for E^-^O^+^ was 0.003 ± 0.002 mV (Fig. 2B), for E^+^O^-^ was 0.01 ± 0.001 mV (Fig. 2C) and for E^+^O^+^L^-^ was 0.001 ± 0.0005 mV (Fig. 2D).

**Figure 2.**
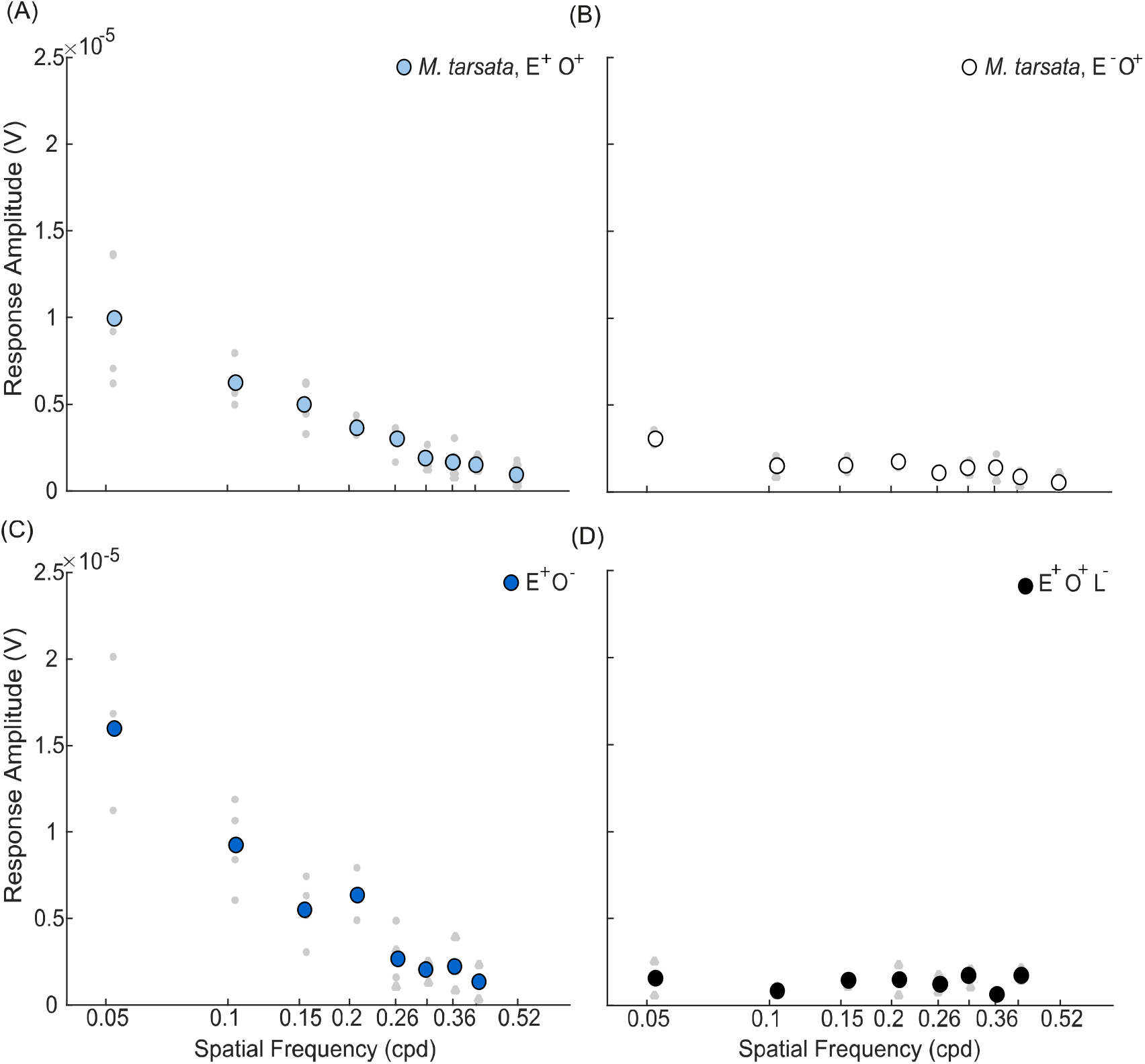
Amplitude of pERG response signal from the ocellar second-order neurons of *Myrmecia tarsata* under different treatment conditions. (**A**) Visual systems-intact individuals (E^+^O^+^), (**B**) Compound eyes-occluded individuals (E^-^O^+^), **(C)** Ocelli-occluded individuals (E^+^O^-^), **(D)** Visual systems-intact individuals with incisions made in the lamina (E^+^O^+^L^-^) (see methods for details). Each coloured data point is the mean amplitude of the signal of all individuals at the corresponding spatial frequency at 95% contrast. Individual data points shown in grey circles. Individual non-significant data points shown in grey triangles. (n=5 for E^+^O^+^, n=4 for E^+^O^-^, n=3 for E^-^O^+^, E^+^O^+^L^-^).

### Spatial properties in visual systems-intact and ocelli-occluded individuals of M. tarsata

We measured contrast sensitivities and the spatial resolving powers in E^+^O^+^ and E^+^O^-^ treated individuals (Fig. 3). At each spatial frequency of the visual stimuli, the amplitude of the pERG response decreased with decreasing contrast. The contrast sensitivity decreased as the spatial frequency increased under both the treatment conditions (Fig. 3A, Table 2). The maximum contrast sensitivities attained at the lowest spatial frequency (0.05cpd) was 12.9 ± 1.2 (mean ± S.E.) in E^+^O^+^ individuals and 6.8 ± 1.2 in E^+^O^-^ treated individuals (Table 1). The spatial resolving power in E^+^O^+^ individuals was 0.29 ± 0.02 cpd (mean ± SE) and 0.21 ± 0.01cpd in E^+^O^-^ treated individuals (Table 1). The contrast sensitivity functions, the maximum contrast sensitivities, and the spatial resolving powers were significantly higher in E^+^O^+^ individuals compared to E^+^O^-^ treated individuals (Tables 2, 3, 4).

**Table 1.**
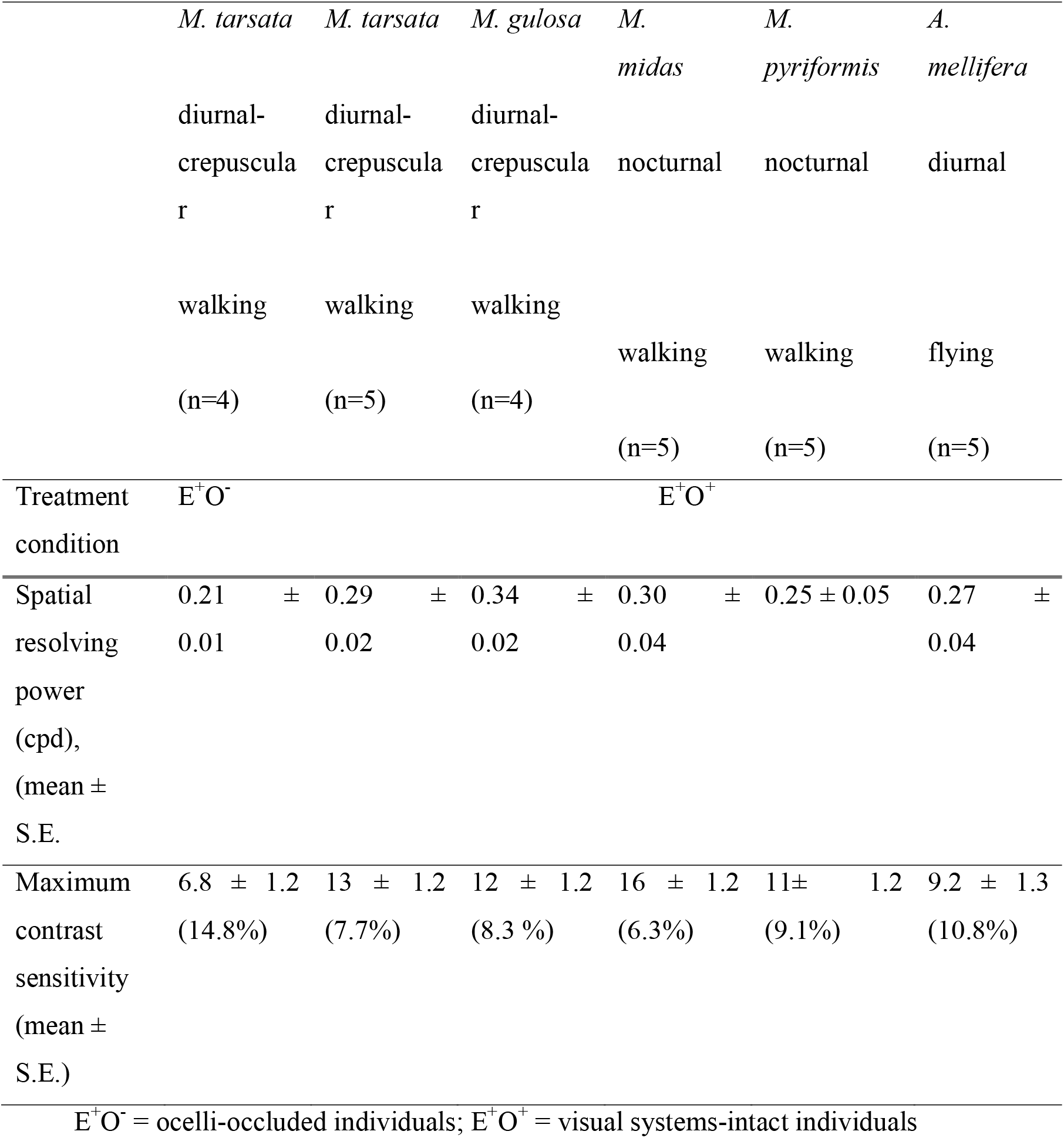
Summary of spatial resolving power and contrast sensitivity of ocellar second-order neurons for *Apis mellifera* and *Myrmecia* ants under different treatment conditions.

**Table 2.** Summary of linear mixed-effects model analysis for testing the effect of spatial frequency of gratings and treatment condition (E^+^O^+^ and E^+^O^-^) on contrast sensitivity functions in *M. tarsata*. Model: contrast sensitivity ∼ spatial frequency + treatment condition (1|animal ID). The *t*-tests for fixed effects use Satterthwaite approximations to degrees of freedom (*df*). The variance of each of the random effects is <3%

| Parameter | Estimate | Standard Error | *df* | *t*-value | *p*-value |
| --- | --- | --- | --- | --- | --- |
| Intercept | -0.5 | 0.09 | 35.74 | -5.3 | <6.03e-06 |
| Spatial frequencies | -1.21 | 0.09 | 31.92 | -12.98 | <2.83e-14 |
| Treatment condition | -0.24 | 0.07 | 7.22 | -3.23 | <0.01 |

**Table 3.** Summary of the linear model for testing the relationship between maximum contrast sensitivity and treatment condition (E^+^O^+^ and E^+^O^-^) in *M. tarsata*. Model: maximum contrast sensitivity ∼ treatment condition

| Parameter | Estimate | Standard Error | *t*-value | *p*-value |
| --- | --- | --- | --- | --- |
| Intercept | 0.1 | 0.07 | 15.25 | 0.000001 |
| Treatment condition | -0.28 | 0.1 | -2.55 | <0.04 |

**Table 4.** Summary of the linear model for testing the relationship between spatial resolving power and treatment condition (E^+^O^+^ and E^+^O^-^) in *M. tarsata*. Model: spatial resolving power ∼ treatment condition

| Parameter | Estimate | Standard Error | *t*-value | *p*-value |
| --- | --- | --- | --- | --- |
| Intercept | 0.29 | 0.02 | 14.87 | <1.49e-06 |
| Treatment condition | -0.08 | 0.03 | -2.75 | <0.03 |

**Figure 3.**
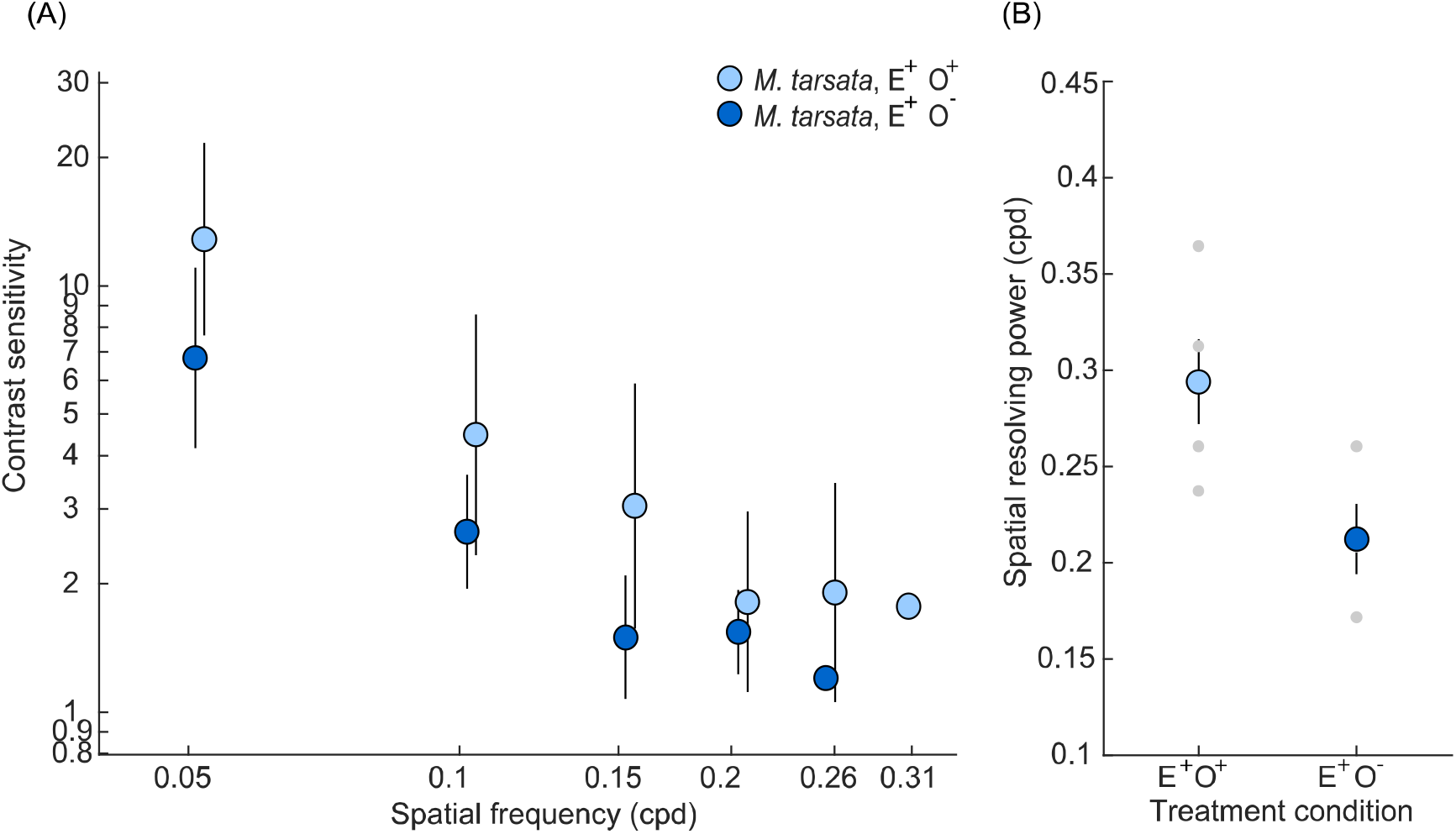
Spatial properties of ocellar second-order neurons of *Myrmecia tarsata* for visual systems-intact individuals (E^+^O^+^) and, ocelli-occluded individuals (E^+^O^-^). **(A) Contrast sensitivity function and (B) Spatial resolving power for the two treatment conditions are shown. In Panel A**, each data point is the mean contrast sensitivity of all individuals of *M. tarsata* for a particular treatment condition at the corresponding spatial frequency. The error bars show 95% confidence intervals. Data points for each condition were shifted to the right or left of the recorded spatial frequency to improve visualisation. **(B)** Each coloured data point is the mean spatial resolving power of all individuals of *M. tarsata* for a particular condition at 95% contrast. Error bars show standard error. Individual data points are shown in grey. (n=5 for E^+^O^+^, n=4 for E^+^O^-^)

### Contrast sensitivity in visual systems-intact individuals of Myrmecia ants and Apis mellifera

We measured the contrast sensitivity of the ocellar second-order neurons in four *Myrmecia* species and in the European honeybee *Apis mellifera* in E^+^O^+^ individuals (Fig. 4). The contrast sensitivity decreased as the spatial frequency increased in all species (Fig. 4A, Table 5). The maximum contrast sensitivities attained at the lowest spatial frequency (0.05cpd) was highest in *M. midas* at 16 ± 1.2 (6.3%) (mean ± S.E.) to the lowest in *A. mellifera* at 9.2 ± 1.3 (10.8%) (Table 1). Among the ants, a linear model of the maximum contrast sensitivities as a function of species showed that the maximum contrast sensitivities did not differ significantly between the species (F_(3,14)_=0.84, *P*=0.49). Additionally, the variation in contrast sensitivity function was explained by the spatial frequency of the gratings, but not by the species or their time of activity (Table 5).

**Table 5.**
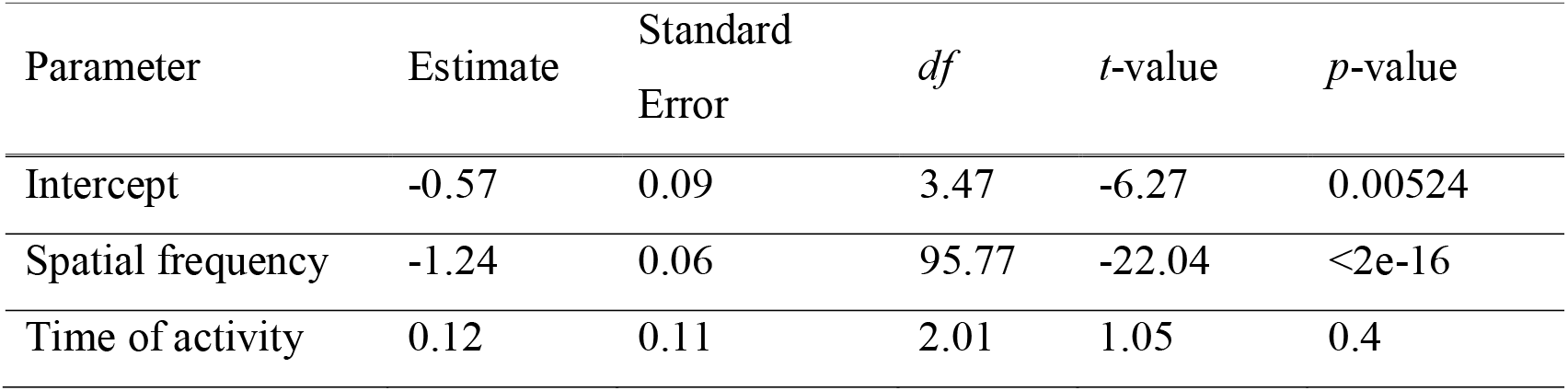
Summary of the linear mixed-effects model for testing the relationship between contrast sensitivity function of the ocellar second-order neurons, spatial frequency of gratings and time of activity in *Myrmecia* ants. Model: contrast sensitivity ∼ spatial frequency + time of activity + (1|species/animal ID). The *t*-tests for fixed effects use Satterthwaite approximations to degrees of freedom (*df*). The variance of each of the random effects is < 3%

**Figure 4.**
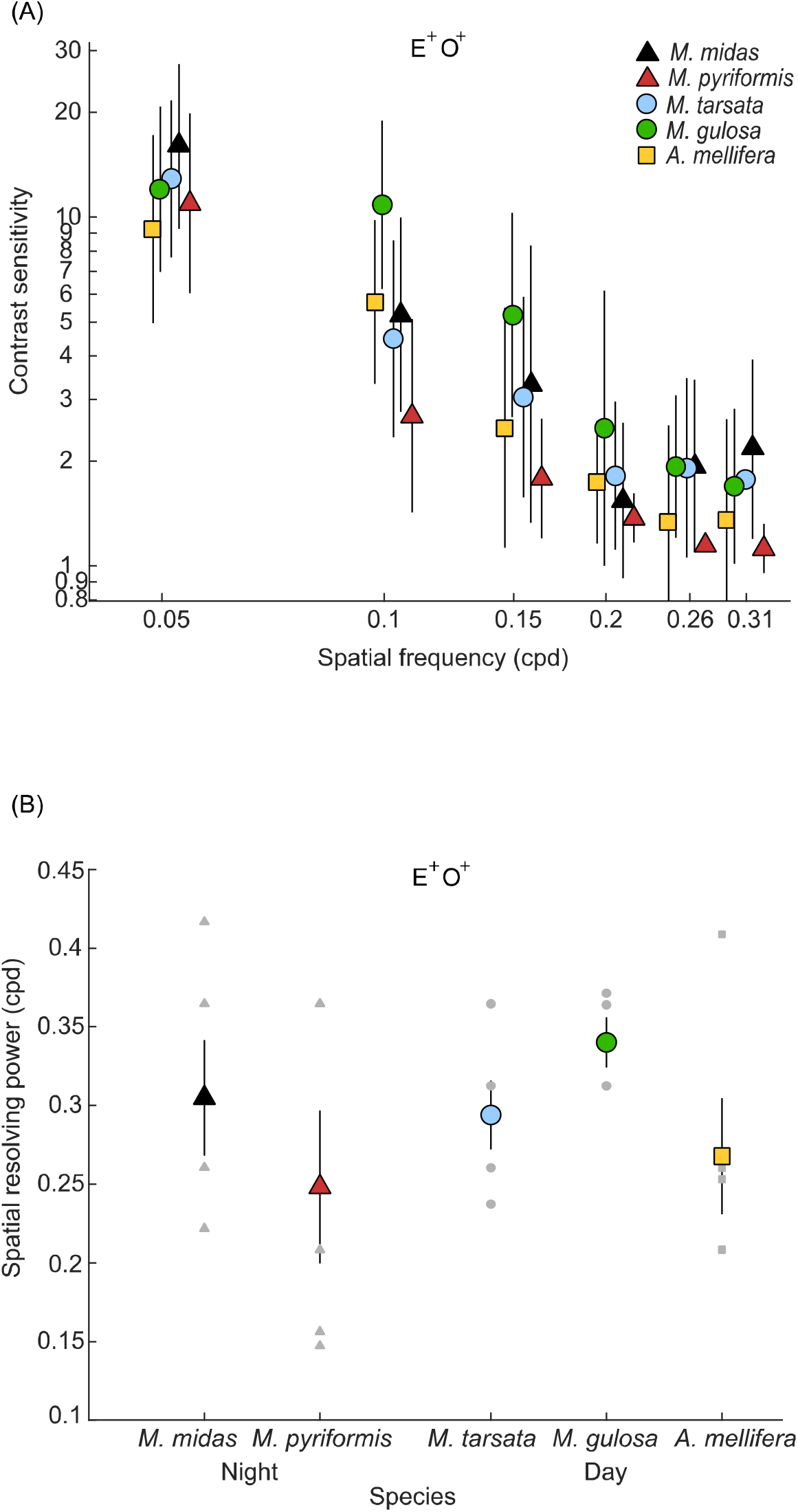
Spatial properties of ocellar second-order neurons of four *Myrmecia* species and *Apis mellifera* for visual systems-intact individuals (E^+^O^+^) **(A) Contrast sensitivity and (B) Spatial resolving power for each species are shown**. In Panel **A**, each data point is the mean contrast sensitivity of all individuals of a particular species at the corresponding spatial frequency. The error bars show 95% confidence intervals. Data points for each species were shifted to the right or left of the recorded spatial frequency to improve visualisation. **(B)** Each coloured data point is the mean spatial resolving power of all individuals of a particular species at 95% contrast. Error bars show standard error. Individual data points are shown in grey. Triangles indicate nocturnal ant species, circles indicate diurnal-crepuscular ant species. (n=4 for *M. gulosa*, n=5 for remaining species)

Due to *Apis mellifera* being a flying diurnal species, we compared its contrast sensitivity function with that of *M. gulosa* and *M. tarsata* which are also active under bright-light conditions but use walking as their primary mode of locomotion. We found that the variation in contrast sensitivity function was explained by the spatial frequency of the gratings, but not by the species or their mode of locomotion (Table 6).

**Table 6.**
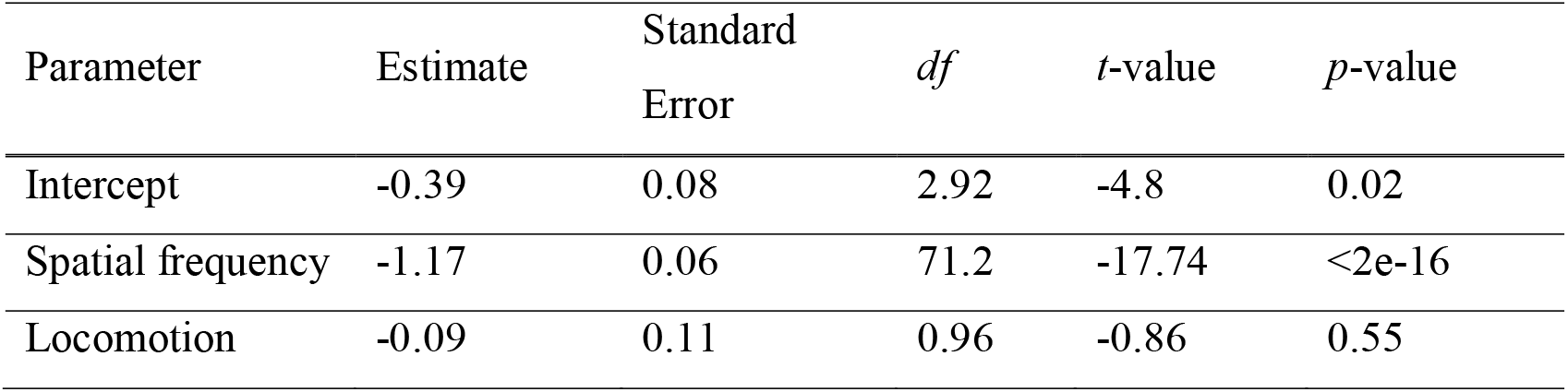
Summary of the linear mixed-effects model for testing the relationship between contrast sensitivity function of the ocellar second-order neurons, spatial frequency of gratings and locomotion in diurnal *Myrmecia* ants and *Apis mellifera*. Model: contrast sensitivity ∼ spatial frequency + locomotion + (1|species/animal ID). The *t*-tests for fixed effects use Satterthwaite approximations to degrees of freedom (*df*). The variance of each of the random effects is < 3%

| Parameter | Estimate | Standard Error | *df* | *t*-value | *p*-value |
| --- | --- | --- | --- | --- | --- |
| Intercept | -0.39 | 0.08 | 2.92 | -4.8 | 0.02 |
| Spatial frequency | -1.17 | 0.06 | 71.2 | -17.74 | <2e-16 |
| Locomotion | -0.09 | 0.11 | 0.96 | -0.86 | 0.55 |

### Spatial resolving power in visual systems-intact individuals of Myrmecia ants and Apis mellifera

The spatial resolving power of the ocellar second-order neurons in the five species was measured in E^+^O^+^ individuals (Fig. 4B). Among the five species, *M. pyriformis* had the lowest spatial resolving power at 0.25 ± 0.05 cpd (mean ± S.E.) and *M. gulosa* had the highest at 0.34 ± 0.02 cpd (Table 1), but this variation was not significantly different between species (Linear model: F_(4,19)_=0.93, *P*=0.47).

## Discussion

### Electroretinogram

To confirm the presence of the extracellular potentials from the ocellar second-order neurons in our ERG recordings, we measured ERGs responding to the ON-OFF stimuli from the ocellar retina of *M. tarsata* for E^+^O^+^ and E^-^O^+^ treated individuals (Fig. 1A). Similar to the ocellar ERGs in other insects (Ruck, 1961a; Ruck, 1961b), we found the presence of a hyperpolarizing post-synaptic potential and a sustained after-potential identified as Component 3 at the stimulus onset (Fig. 1B) and offset (Fig. 1C) respectively for E^+^O^+^ individuals. In E^-^O^+^ treated individuals, the sustained after-potential identified as part of Component 3 was seen at the stimulus offset (Fig. 1C). This confirms the presence of ERGs from the ocellar second-order neurons in our pERG recordings. As mentioned earlier, some L-neurons are known to interact with descending interneurons which receive input from the compound eyes (e.g., Strausfeld, 1976). We suspect that the presence of these descending interneurons along with the ocellar second-order neurons may explain the presence of Component 3 at both offset and onset in E^+^O^+^ individuals. However, when the compound eyes are occluded (E^-^O^+^), the ocellar second-order neurons alone produce Component 3 at the stimulus offset, although the Component 3 is less pronounced at the stimulus onset.

### Contribution of the two visual systems to the pERG response amplitude

In order to evaluate the contribution of the signals from the compound eyes and the ocelli to the ocellar second-order neurons, we measured the pERG response amplitudes in E^+^O^+^, E^+^O^-^, E^-^O^+^ and E^+^O^+^L^-^ treated individuals of *M. tarsata* (Fig. 2). At the lowest spatial frequency (0.05cpd) although the mean response amplitude was higher in E^+^O^-^ individuals (Fig. 2C) than in E^+^O^+^ individuals (Fig. 2A), the response amplitudes were not significantly different. This indicates that the compound eyes on their own are adequate to generate a significant pERG response amplitude. The mean response amplitudes at the lowest spatial frequency (0.05 cpd) were higher in conditions when the compound eyes were intact (Fig. 2A, 2C) compared to conditions when the compound eyes were occluded or their neural input disrupted (Fig. 2B, 2D). This indicates that the compound eyes contribute highly to the response amplitude of the ocellar second-order neurons.

The compound eyes’ contribution onto ocellar second-order neurons in ants may be explained by the neural pathways discussed previously. In various insect species, certain L-neurons are known to interact with descending interneurons (e.g., Strausfeld, 1976). We suspect a similar pathway is present in the ants tested. Additionally, the scale of the contribution of the compound eyes (Fig. 2A, C) was unexpectedly much higher compared to the contribution of the ocelli to the ocellar second-order neurons (Fig. 2B). This may simply be a consequence of the high light capturing ability of the compound eyes owing to its multi-lens structure with a large number of facets (*M. tarsata*: 2724 ± 67 facets/eye) (Greiner et al., 2007) and therefore numerous rhabdoms. This is drastically different when compared to the ocelli that consists of a single but large lens and has fewer rhabdoms (*M. tarsata*: 46.5 ± 7 rhabdoms in the median ocelli) (Narendra and Ribi, 2017). Overall, insects with both visual systems appear to largely use their compound eyes to obtain sufficient spatial resolution and sensitivity in order to perform various visually guided behaviours.

Incidentally, the mean response amplitude at the lowest spatial frequency (0.05cpd) was significantly higher in E^-^O^+^ individuals (Fig. 2B) than in E^+^O^+^L^-^ individuals (Fig. 2D). These differences may be due to the disruption in the neural pathway in E^+^O^+^L^-^ treated individuals. Consequently, this method was more effective to block the compound eyes’ input than the compound eyes’ occlusion done using black paint in the E^-^O^+^ individuals.

### Ocellar contrast sensitivity in M. tarsata

Due to the extremely low pERG response amplitudes from the ocellar second-order neurons of the E^-^O^+^ treated individuals (Fig. 2B), it was not possible to directly measure the contribution of the ocelli to the contrast sensitivity and the spatial resolving power of the ocellar second-order neurons in *M. tarsata*. However, the high pERG response amplitudes in the E^+^O^+^ treated individuals (Fig. 2A) and E^+^O^-^ treated individuals (Fig. 2C) enabled us to estimate the visual capabilities of the ocelli. We found that the contrast sensitivity function and the maximum contrast sensitivity of ocellar second-order neurons were significantly different in E^+^O^+^ and E^+^O^-^ treated individuals. The mean maximum contrast sensitivity was higher in E^+^O^+^ treated individuals (13 ± 1.2 (7.7%), mean ± SE) than that in E^+^O^-^ treated individuals (6.8 ± 1.2 (14.8%)), indicating a significant contribution of the ocelli to contrast sensitivity of the ocellar second-order neurons. This demonstrates that inputs from both the ocelli and the compound eyes contribute to the contrast sensitivity of the ocellar second-order neurons.

We speculate that in ants the descending interneurons receive information from the ocelli first and subsequently from the compound eyes. This is quite possible due to the fast transmission of signals via the L-neurons found in several insects (Guy et al., 1979; Mizunami, 1995). This could either modulate or gate the signals from the compound eyes (Guy et al., 1979). The input from the compound eyes could further increase the contrast sensitivity of the ocelli to enable efficient navigation (eg., Fent and Wehner, 1985b; Schwarz et al., 2011b). Increased contrast sensitivity of the ocellar second-order neurons based on the contribution of the compound eyes together with the fast transmission of signals through the L-neurons would be beneficial for navigation and other visually guided behaviours (eg., Barry and Jander, 1968; Cornwell, 1955; Honkanen et al., 2018; Lindauer and Schricker, 1963; Schricker, 1965; Wellington, 1974)

### Ocellar spatial resolving power in M. tarsata

The spatial resolving power of the ocellar second-order neurons in E^+^O^+^ individuals (0.29 ± 0.02 cpd) and E^+^O^-^ treated individuals (0.21 ± 0.01 cpd, Fig. 3B) were significantly different. Although a large number of ocellar photoreceptors converge onto very few ocellar second-order neurons (Berry et al., 2006; Chappell et al., 1978; Goodman and Williams, 1976; Guy et al., 1979; Mizunami, 1995; Patterson and Chappell, 1980; Toh and Kuwabara, 1974), in dragonflies the spatial resolution is conserved even after this convergence indicating the possibility of local processing within the ocellar neuropil (Berry et al., 2006). Our results suggest that in ants, the ocellar second-order neurons are involved in processing such that they contribute to ocellar spatial vision to some extent but its functional significance remains to be investigated. It is likely that the ocellar spatial acuity is enhanced when there is input from the both the ocelli and compound eyes.

### Spatial properties of the ocellar second-order neurons in ants and honeybees

Nocturnal *Myrmecia* ants as previously described have larger ocelli and wider ocellar rhabdoms (Narendra and Ribi, 2017) compared to their diurnal relatives indicating that the ocelli might have higher contrast sensitivities. Therefore, we compared the contrast sensitivities of the ocellar second-order neurons in E^+^O^+^ individuals of *Myrmecia* species active at discrete times of the day (Fig. 4). However, the contrast sensitivity functions were not significantly different between the species indicating that the time of activity did not have an effect on their contrast sensitivity functions. This is consistent with our knowledge on the compound eyes where the time of activity did not explain the difference in contrast sensitivities of the compound eyes of diurnal and nocturnal *Myrmecia* ants (Ogawa et al., 2019).

To identify whether ocellar spatial properties were affected by the mode of locomotion, we studied the diurnal flying European honeybee *A. mellifera* (E^+^O^+^) and compared it with diurnal *Myrmecia* ants. We chose *A. mellifera* because their ocelli has been well studied: interaction with the compound eyes has been mapped (Guy et al., 1979); the plane of best focus is known to lie on the ocellar retina (Ribi et al., 2011). With ocellar lens diameters of 294 µm *Apis mellifera* (Ribi et al., 2011)) have distinctly larger ocelli than the *Myrmecia* ants (*M. tarsata*: 129.2 µm (Narendra and Ribi, 2017)). Our results showed that the species and their locomotion did not have an effect on their contrast sensitivity functions.

Based on our experimental paradigm, we suspect that the main function of the ocellar second-order neurons for E^+^O^+^ individuals in all five species is to detect overall bright and dim contrasts indicating that the overall processing of visual information is similar in their peripheral neural pathways. However, it must be stated that the contrast sensitivities of species may change depending on the intensity of light present. Our results are reflective of contrast sensitivities at a particular light intensity and may differ at different light intensities.

The spatial resolving powers of the ocellar second-order neurons of *Myrmecia* ants and that of *Apis mellifera* for E^+^O^+^ individuals measured in this study were not significantly different between species. The neural pathways relaying information from the compound eyes to the ocellar second-order neurons are likely to be conserved across all the five Hymenopteran species leading to the lack of difference in spatial resolving power in all species.

In conclusion, our results provide physiological evidence that the compound eyes mediate the signals from the ocelli therefore significantly affecting the contrast sensitivity of the median ocelli and subsequently the spatial resolving power with a noticeably larger improvement in contrast sensitivity than the spatial resolving power. The consequence of the compound eyes modulating the signals from the ocelli and how this information is processed in the higher order brain regions and translated into visually guided behaviours remains to be discovered.

## Competing interests

The authors declare no competing or financial interests.

## Author contributions

Study design: B.P., Y.O., A.N.; Data collection and analyses: B.P., Y.O., L.R.; Built the equipment and wrote the software: N.H., L.R., Y.O.; Writing – original draft: B.P.; Writing – review and editing: all authors; Visualisation: B.P., Y.O., A.N.; Funding acquisition: A.N.

## Funding

We acknowledge financial support from the Australian Research Council, Discovery Project grants (DP150101172, DP200102337) and Future Fellowship (FT140100221). B.P. was supported by the International Macquarie Research Excellence Scholarship.

## Notes

### Competing Interest Statement

The authors have declared no competing interest.

